# Differences in codon usage between host-species-specific rabies virus clades are driven by non-GC3 nucleotide composition

**DOI:** 10.64898/2026.01.12.699068

**Authors:** Rowan Durrant, Jonathan Dushoff, Matthew Arnold, Christina Cobbold, Katie Hampson

## Abstract

Viral genes sometimes use certain codons more than others due to their nucleotide content, translational efficiency, and selection pressure from the host immune system. The rabies virus (RABV) is a negative strand RNA virus which can infect a broad range of mammalian hosts, with many of its clades circulating predominantly in specific host species. Previous work on codon usage in RABV has focused only on broader viral clades. We use publicly available RABV nucleoprotein gene sequences to investigate how dinucleotide content and codon usage biases differ between host-associated clades, and what drives these differences. We found that codon usage varies most between bat- and carnivore-associated RABV clades, and more subtly between host-species-specific minor clades within these groups. C1, GA3 and GT3 content were found to have a strong influence over codon usage patterns, and CpG content was higher in carnivore-associated RABV clades than in bat-associated clades. This, along with lower numbers of zinc-finger antiviral protein binding motifs than would be expected based on (di)nucleotide composition in bat-associated RABV sequences, suggests that bat-associated RABV clades may be under higher selection pressure from the host’s zinc-finger antiviral protein than carnivore-associated clades are, warranting further investigation of the mechanism underpinning this change.

## Introduction

Despite encoding the same amino acid, synonymous codons are often not used equally. While a wide range of factors likely influence these imbalances (Parvathy *et al*. 2022), codon usage biases in both RNA and DNA viruses generally appear to be shaped by nucleotide-level effects such as guanine and cytosine (GC) content rather than by codon-level translational selection (Jenkins & Holmes 2003; Shackelton *et al*. 2006). Unlike positive-strand RNA viruses (Tian *et al*. 2018), negative-strand RNA viruses do not appear to mimic their host species’ codon usage patterns (Rima 2015), despite also being reliant on the host’s translation machinery. This may be due to transcription of viral RNA being dependent on RNA stability, which is influenced by codon usage (Gumpper *et al*. 2019). Nonetheless, using machine learning methods, codon usage together with nucleotide and dinucleotide content can accurately predict the hosts of coronaviruses (Brierley & Fowler 2021) and other RNA viruses (both positive-and negative-strand; Babayan *et al*. 2018), suggesting a connection between viral codon usage and host species.

The rabies virus (RABV) is a negative-strand RNA virus that can infect a broad range of mammal species. The RABV genome consists of five genes, which encode the nucleoprotein (N), phosphoprotein (P), matrix protein (M), glycoprotein (G) and RNA polymerase (L; “large” protein). RABV can be broadly split into bat- and carnivore-associated clades, with the main bat-to-carnivore host shift event being estimated to have occurred approximately 600 years ago (Troupin *et al*. 2016). Another shift, from bats to skunks and raccoons, is estimated to have occurred 250 years ago (Ding *et al*. 2017), and bat-to-carnivore spillover events continue to occur in the present day (Kuzmin *et al*. 2012). Some RABV clades appear to be highly host-specific, circulating within a single species; for example, sustained transmission cycles have been observed in Chinese ferret badger (CFB) populations (Zhang *et al*. 2013) associated with the minor clade Asian SEA2b, as opposed to the clade circulating sympatrically in dogs, Asian SEA2a.

There has been debate as to whether these host-species-specific transmission cycles are due to adaptation or ecological circumstances (Marston *et al*. 2018; Mollentze *et al*. 2014), with evidence of positive selection over host shifts being mixed: some studies show no evidence of adaptation to new host species (Kuzmin *et al*. 2012; Troupin *et al*. 2016), while others observed some positively selected sites but with low repetition across different host shift events (Streicker *et al*. 2012). Troupin *et al*. (2016) also found, however, that an amino acid change from leucine to serine at nucleoprotein position 374 appears ubiquitously, and almost exclusively, in two separately emerging ferret badger-associated clades, which suggests that this amino acid change is required for sustained transmission in ferret badgers. These studies focused on non-synonymous substitutions (changes to the amino acid sequence and therefore the structure of the encoded protein), leaving the possible influence of synonymous substitutions (i.e., changes in codon usage bias or dinucleotide bias) on RABV-host adaptation yet to be investigated.

On average, across all available whole genome sequences, RABV has the most balanced codon usage among lyssaviruses (Zhang *et al*. 2018), perhaps due to its broad host range compared to other members of the family (Marston *et al*. 2018). Despite this, codon selection is important to RABV fitness; viral replication is inhibited when the RABV matrix protein sequence is reconstructed with codons that are rarely used in the cells that the virus is cultured in (Luo *et al*. 2020a). This may be due to a collateral increase in CpG and UpA dinucleotides, which are selected against by host immune mechanisms such as the zinc-finger antiviral protein (ZAP; Tulloch *et al*. 2014). Previous codon usage studies focusing on the rabies virus either did not investigate differences in codon usage on a clade level (Morla *et al*. 2016), or subdivided the sequences into the major RABV clades (Bat, Africa-2, Arctic, Cosmopolitan, Asian and Indian subcontinent major clades in He *et al*. (2017); plus Africa-3 and South-East Asian clades in Li *et al*. (2023), potentially losing signal from host-species-specific minor clades. Investigating differences in codon usage between host-species-specific minor clades has the potential to shed light on the effect of sustained transmission within a single host species on codon usage.

In this study we aimed to quantify differences in codon usage biases between host-species-specific RABV minor clades, and investigate what factors underpin these differences. We hypothesised that clades that diverged more deeply in the virus’s evolutionary history (i.e., from bats to carnivores) would show clear codon usage biases due to longer periods of sustained transmission within specific host groups, while more recent host shifts (i.e., to Chinese ferret badgers) would show less obvious differences to their most closely related clades.

## Methods

### Data acquisition

We downloaded complete RABV N gene sequences from the RABV-GLUE sequence database (Campbell *et al*. 2022; acquired 23 March 2026). The N gene was chosen as it was the most commonly sequenced gene across a wide range of clades. We searched for sequences with 100% N gene coverage from a selection of host-species-specific minor clades spanning a wide range of host species, including those mentioned by (Troupin *et al*. 2016) and four bat-associated clades. The full list of clades and the terms used to filter sequences on RABV-GLUE are shown in table 1.

**Table 1:** RABV-GLUE clades and filtering terms used when downloading RABV N gene sequences. Date ranges include only sequences used in the final dataset.

| Clade | Host | RABV-GLUE host filter term: contains | No. sequences (initial → deduplicated → no ambiguous sites) | Sampling date range |
| --- | --- | --- | --- | --- |
| Asian SEA2a | Domestic dog | “Canis familiaris”, “Canis lupus familiaris” | 77 → 59 → 59 | 1994-2016 |
| Asian SEA2b | Chinese ferret badger | “Melogale”, “ferret badger” | 50 → 29 → 29 | 2008-2017 |
| Arctic A | Arctic fox | “Vulpes lagopus”, “Alopex”, “arctic fox” | 104 → 72 → 69 | 1977-2017 |
| Cosmopolitan AM2a | Mongoose | “Mongoose” | 9 → 9 → 9 | 2000-2002 |
| Cosmopolitan AF1b | Domestic dog | “Canis familiaris”, “Canis lupus familiaris” | 232 → 129 → 116 | 1986-2022 |
| RAC-SK SCSK | Skunk | “Mephitis”, “skunk” | 9 → 9 → 9 | 2009-2017 |
| Bat DR | Vampire bat | “Desmodus rotundus” | 56 → 41 → 40 | 1988-2024 |
| Bat TB1 | Mexican free-tailed bat | “Tadarida brasiliensis” | 32 → 31 → 31 | 2002-2005 |
| Bat EF-E2 | Big brown bat | “Eptesicus fuscus” | 36 → 35 → 35 | 2003-2005 |
| Bat LC | Hoary bat | “Lasiurus cinereus” | 17 → 17 → 16 | 1998-2005 |

Both raccoon- and skunk-associated RAC-SK sequences were initially included in the dataset, but the majority of these sequences were removed after initial phylogenetic analysis suggested substantial transmission between raccoons and skunks within this clade. The southern central skunk variant (SCSK) within the RAC-SK major clade did not contain any raccoon-derived sequences, and was therefore retained. We downloaded 622 N gene sequence alignments, constrained to RABV-GLUE’s reference N genes with a length of 1353 nucleotides, directly from RABV-GLUE. No internal stop codons or gap regions were observed. Duplicate sequences (N = 191) and sequences containing ambiguous bases (i.e., N, M, W, R or Y; N = 18) were removed from the dataset, leaving 413 complete N gene sequences in total. The GenBank accession numbers for these sequences are available in Supplementary Table S1.

### Phylogenetic trees

Phylogenetic trees were constructed using IQ-TREE version 1.6.12 (Nguyen *et al*. 2015) with ultrafast bootstrapping with 1000 replicates, and the best-fitting model determined by IQTree’s ModelFinder (GTR+F+R3). European Bat Lyssavirus 1 (Accession no. NC_009527.1), Gannoruwa Bat Lyssavirus (NC_031988) and Australian Bat Lyssavirus (NC_003243.1) sequences were used as outgroups; using only a single lyssavirus sequence as an outgroup resulted in an unresolved root node. Sequences were aligned using MAFFT (Katoh *et al*. 2002). Visualisation was carried out in R version 4.5.1 (R Core Team 2025) using the *ggtree* R package (Yu *et al*. 2017).

### Codon usage

We used the effective number of codons (ENC; a measure of overall codon usage bias accounting for amino acid composition, with values ranging from 20 to 61, where lower values correspond to stricter biases) to compare the strength of codon usage biases between the RABV clades, calculated using the equations shown in (Wright 1990):

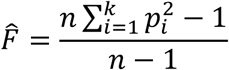

where 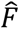 is the homozygosity of each amino acid, 𝑝_𝑖_ is the frequency of the synonymous codon *i*, *k* is the number of codons that encode the amino acid, and *n* is the total frequency of the amino acid; and:

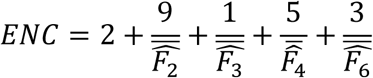

where 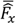 is the mean homozygosity of the group of amino acids encoded by *x* synonymous codons.

Raw codon usage and relative synonymous codon usage (RSCU) values were calculated for each minor clade using CAIcal (Puigbò *et al*. 2008), which uses the following formula (Sharp & Li 1986):

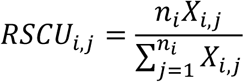

where 𝑛_𝑖_ is the number of synonymous codons for amino acid *j*, and 𝑋_𝑖,𝑗_ is the number of occurrences of codon *i* for amino acid *j*. The raw codon usage counts (excluding amino acids with only one synonymous codon) were analysed using principal component analysis (PCA) using the *stats* R package (R Core Team 2025), and the correlation coefficients between the resulting PC values and 46 nucleotide composition metrics (nucleotide and nucleotide pair composition overall and at each codon position, obs/exp CpG content and counts, and obs/exp UpA content and counts) were calculated. We then used a phylogenetic mixed model to account for genetic relatedness between the N gene sequences when comparing the PC values to their most highly correlated metric using the *MCMCglmm* R package (Hadfield 2010) with the default normal distribution prior. Chains were run for 15,000 iterations with 10% discarded as burn-in to ensure an effective sample size above 1,000, and traces were visually inspected for convergence. We also compared the PC loadings to the composition metrics of each of the 59 codons used in the PCA and repeated the PCA with RSCU values instead of raw codon counts to confirm whether these results were codon-level effects, or whether they were influenced by differences in amino acid content.

Codon adaptation index (CAI) values were calculated for each RABV clade against four host species using CAIcal to measure the similarity in codon usage between each RABV clade and the corresponding host species. CAI is calculated from a viral sequence and a host reference codon usage table with the following equation (Sharp & Li 1987):

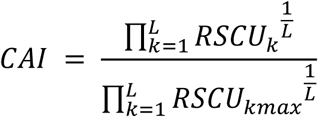

where for each codon k in the viral sequence, 𝑅𝑆𝐶𝑈_𝑘_ is the RSCU of codon *k* in the host reference table, 𝑅𝑆𝐶𝑈_𝑘𝑚𝑎𝑥_ is the maximum RSCU in the host reference table for the amino acid encoded by codon *k*, and *L* is the total number of codons in the viral sequence (excluding the start codon). Host genomic codon usage datasets were obtained for domestic dogs (*Canis familiaris*), Arctic foxes (*Vulpes lagopus*), big brown bats (*Eptesicus fuscus*) and vampire bats (*Desmodus rotundus*) from CoCoPUTs’ RefSeq dataset (Alexaki *et al*. 2019). Other host species had no available codon usage data, so all 10 RABV clades in our dataset were compared to the above four host species. The CAI values were divided by the expected CAI (eCAI; a 95% upper confidence limit for CAI of a random sequence, controlling for amino acid usage and GC content (Puigbò *et al*. 2008)) values to give a normalised CAI (nCAI). We investigated the relationship between nCAI and UpA or CpG content using *MCMCglmm* to account for genetic relatedness, with the default normal distribution prior. Chains were run for 15,000 iterations with 10% discarded as burn-in.

### Dinucleotide content

The ratio of observed-to-expected CpG and UpA content for each sequence was calculated as in (Gardiner-Garden & Frommer 1987):

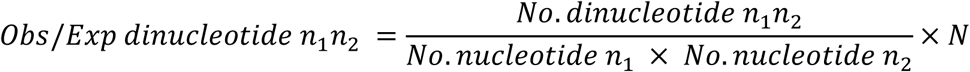

where *N* is the nucleotide length of the sequence, and *n*_1_ and *n*_2_ are the first and second nucleotides in the dinucleotide respectively. Differences in dinucleotide content between groups of clades infecting bats or carnivores (i.e., the clades Bat LC, EF-E2, DR and TB in the bat-associated clades group, versus all other clades including RAC-SK SCSK in the carnivore-associated clades group) were assessed while accounting for phylogeny using the *MCMCglmm* R package with the default normal distribution prior (Hadfield 2010). Chains were run for 15,000 iterations with 10% discarded as burn-in.

Marginal ancestral sequence reconstruction was built from the IQ-TREE trees described above. The ancestral sequence at each node was reconstructed 100 times by sampling the reported probabilities of each nucleotide at each site. The ratios of observed-to-expected CpG and UpA content of the resulting ancestral sequences were then calculated to examine when in the rabies virus’s evolutionary history changes in CpG and UpA content may have occurred.

We also compared the frequency of ZAP’s CpG-based optimal binding motif, C(n_m_)G(n)CG (where n can be any nucleotide, and the subscript m denotes the length of this subsequence, being either 4, 5, 6, 7 or 8 nucleotides long, determined in mice (Luo *et al*. 2020b)), to the expected number of these motifs for each sequence given their nucleotide and dinucleotide contents. This expected value was generated by reshuffling each RABV sequence 100 times using the *uShuffleR* R package (which is based on the uShuffle algorithm; Jiang *et al*. 2008), maintaining the nucleotide and dinucleotide content of the original sequence, and calculating the mean number of ZAP binding motifs in these reshuffled sequences. The CpG-based binding motif has been previously analysed in SARS-CoV-2 (Afrasiabi *et al*. 2022) and hepatitis C virus (Mukherjee *et al*. 2023) sequences, with results suggesting that the binding motif found in mice is shared by humans; little is known about ZAP binding motifs in other mammalian species. The UpA-based ZAP binding motif is currently unknown, so we could not perform a corresponding UpA-based investigation.

## Results

The RABV phylogeny can be broadly split into carnivore-associated (Cosmopolitan AF1b, Cosmopolitan AM2a, Arctic A, Asian SEA2a and Asian SEA2b) and bat-associated (Bat DR, Bat TB1, Bat LC and Bat EF-E2) clades, with the exception of the skunk-associated RAC-SK SCSK clade, which groups with the bat-associated clades (Figure 1). This clade is thought to have descended from bat-associated clades separately to the main bat-to-carnivore host shift (Ding *et al*. 2017; Troupin *et al*. 2016). The only paraphyletic RABV-GLUE-designated clade was the Asian SEA2a domestic dog-associated clade, which also contains the more recently diverged Asian SEA2b clade, associated with Chinese ferret badgers.

**Figure 1:**
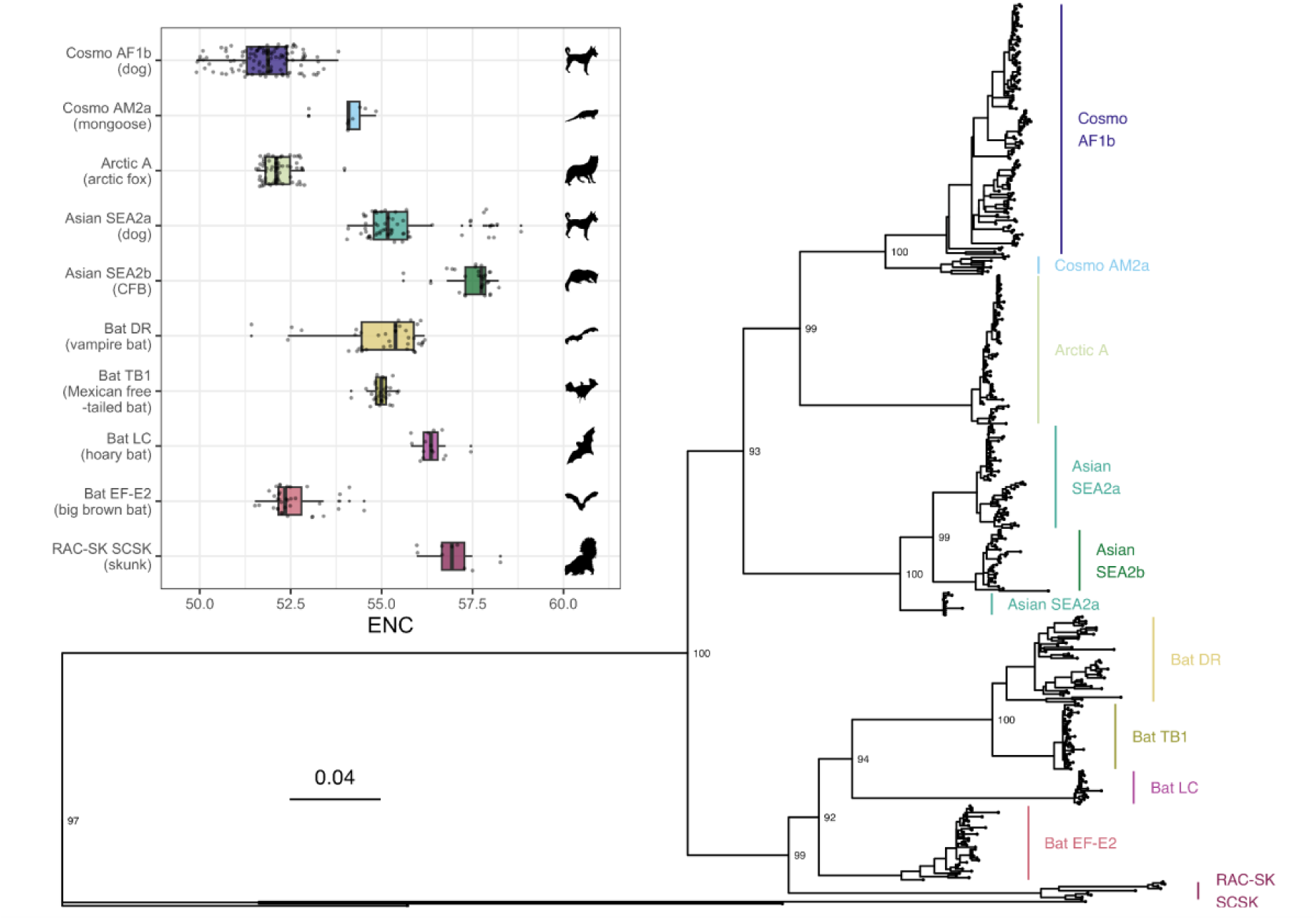
Maximum-likelihood phylogenetic tree of all the sequences used in analyses. Three non-rabies lyssavirus sequences were used as an outgroup. Branch lengths are scaled by substitutions per site. Internal node labels are bootstrap support values. **Inset:** Effective number of codons (ENC) across different host-species-specific rabies clades. Points are coloured by species-specific minor clade. An ENC of 61 denotes no tendency to prefer one synonymous codon over another, whereas an ENC of 20 denotes strict use of one codon per amino acid. CFB = Chinese ferret badger.

### Codon usage

The effective number of codons (ENC) is a measure of bias in codon usage, where a value of 61 denotes that each of the 61 codons are represented equally within their corresponding amino acid, and a value of 20 means only one codon is used to encode each of the 20 amino acids. The mean ENC used by the rabies virus N gene was calculated to be 53.7 (95%CI: 53.5, 54.0) with a standard deviation of 2.12, suggesting little bias in codon usage, but this varied slightly between host-specific minor clades (Figure 1 inset). The dog-associated Cosmopolitan AF1b clade was the most biased in its codon usage (mean ENC of 51.8; 95%CI: 51.6, 52.0), with the big brown bat and arctic fox-associated clades (Bat EF-E2 and Arctic A, respectively) also being more biased than other clades (mean ENCs of 52.6 (95%CI: 52.3, 52.8) and 52.2 (95%CI: 52.1, 52.3) respectively). The Chinese ferret badger-associated clade Asian SEA2b was the least biased (mean ENC of 57.5; 95%CI: 57.3, 57.7).

Certain amino acids appear to have stronger codon biases than others (Figure 2; Supplementary Table S2); the arginine (R) codon AGA has an RSCU value above 2 in every host-species-specific clade investigated (RSCU values above 1.6 suggest strong overrepresentation), and was the only codon that was consistently preferred in all of the host-species-specific minor clades investigated. For every other amino acid, the most commonly preferred codon varied between clades. Only four amino acids (isoleucine, asparagine, aspartic acid and glutamic acid) had no codons that were strongly over- or underrepresented (RSCU above 1.6 or below 0.6) in any clade. While preference for codons containing a UpA dinucleotide varied between codons and clades, codons containing a CpG dinucleotide were almost universally underrepresented.

**Figure 2:**
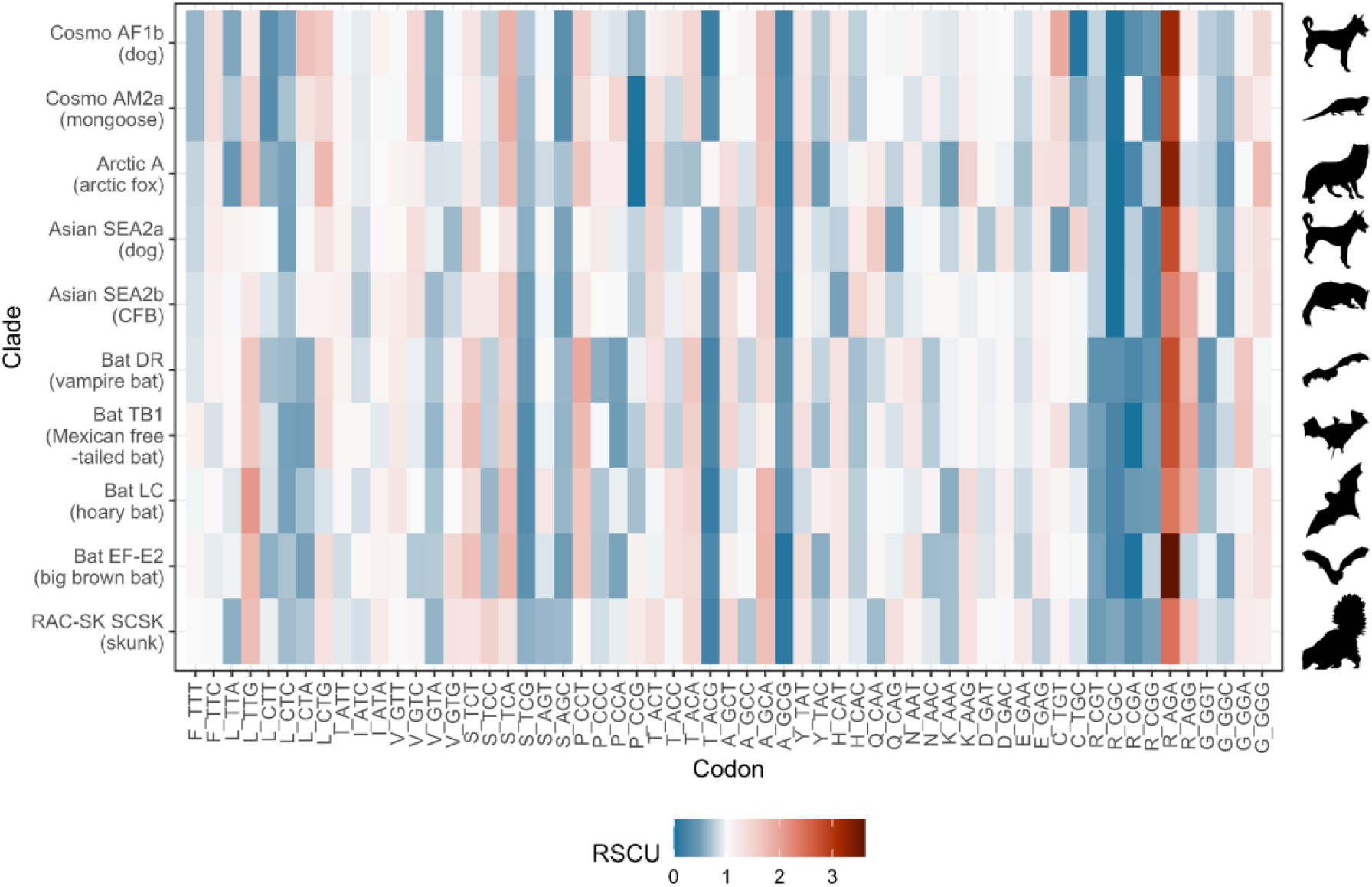
Strength of codon preference varies by amino acid. Cell colour represents RSCU value, where white indicates no codon preference, red indicates preference towards the codon, and blue preference against. CFB = Chinese ferret badger.

We then conducted a principal component analysis (PCA) on the raw codon usage values to further explore any patterns in codon usage between clades and investigate what factors influence these differences. The first and second principal components accounted for 22.4% and 20.6% of the total variation in codon usage respectively, and PC3 accounts for an additional 16.5% of variation (Figure 3A-C; Supplementary Figure S1). Some host-specific RABV clades appear to group by their major clade, with bat-associated and carnivore-associated clades broadly splitting along PC1 (excepting Cosmopolitan AF1b and RAC-SK SCSK), and Cosmopolitan and Asian clades splitting along PC2. PC1 values are correlated with cytosine content at codon position 1 (adjusted R^2^ of 0.666 (95%CI 0.620 - 0.708 *via* linear model bootstrapping with 10,000 replicates); Figure 3D), and this relationship is significant when phylogenetic structure is accounted for using MCMCglmm (pMCMC < 0.001). PC2 values are strongly correlated with the purine content at the third codon position (adjusted R^2^ of 0.729 (95%CI 0.688 - 0.768); pMCMC < 0.001; Figure 3E), and PC3 is very strongly correlated with guanine and thymine content at the third codon position (adjusted R^2^ of 0.840 (95%CI 0.814 - 0.864); pMCMC <0.001; Figure 3F). The loadings of each codon for PC1, PC2 and PC3 are given in Supplementary Table S3. While the GT3 content of the individual codons has a moderate correlation with the PC3 loadings (r = -0.425; 95%CI: 0.189 - 0.614 *via* bootstrapping), suggesting that this relationship represents a codon-level effect, the correlation between the codon’s C1 content and the PC1 loadings and GA3 content and the PC2 loadings is weak (r = -0.196 (95%CI: -0.430 - 0.064) and 0.110 (95%CI: -0.150 - 0.356) respectively; Figure 3G), suggesting that these principal components may be influenced by amino acid composition to some extent. Repeating the PCA using RSCU values instead of raw codon counts gives similar clusters, but with some differences in PC1 and PC2 (Supplementary Figure S2), supporting the presence of some amino acid-level clustering in the raw codon count PCA. The points of the ENC-GC3 plot do not follow the curve that would be expected if codon usage biases are influenced purely by GC content, indicating that this is not a major driver of codon usage bias in RABV (Figure 3H).

**Figure 3:**
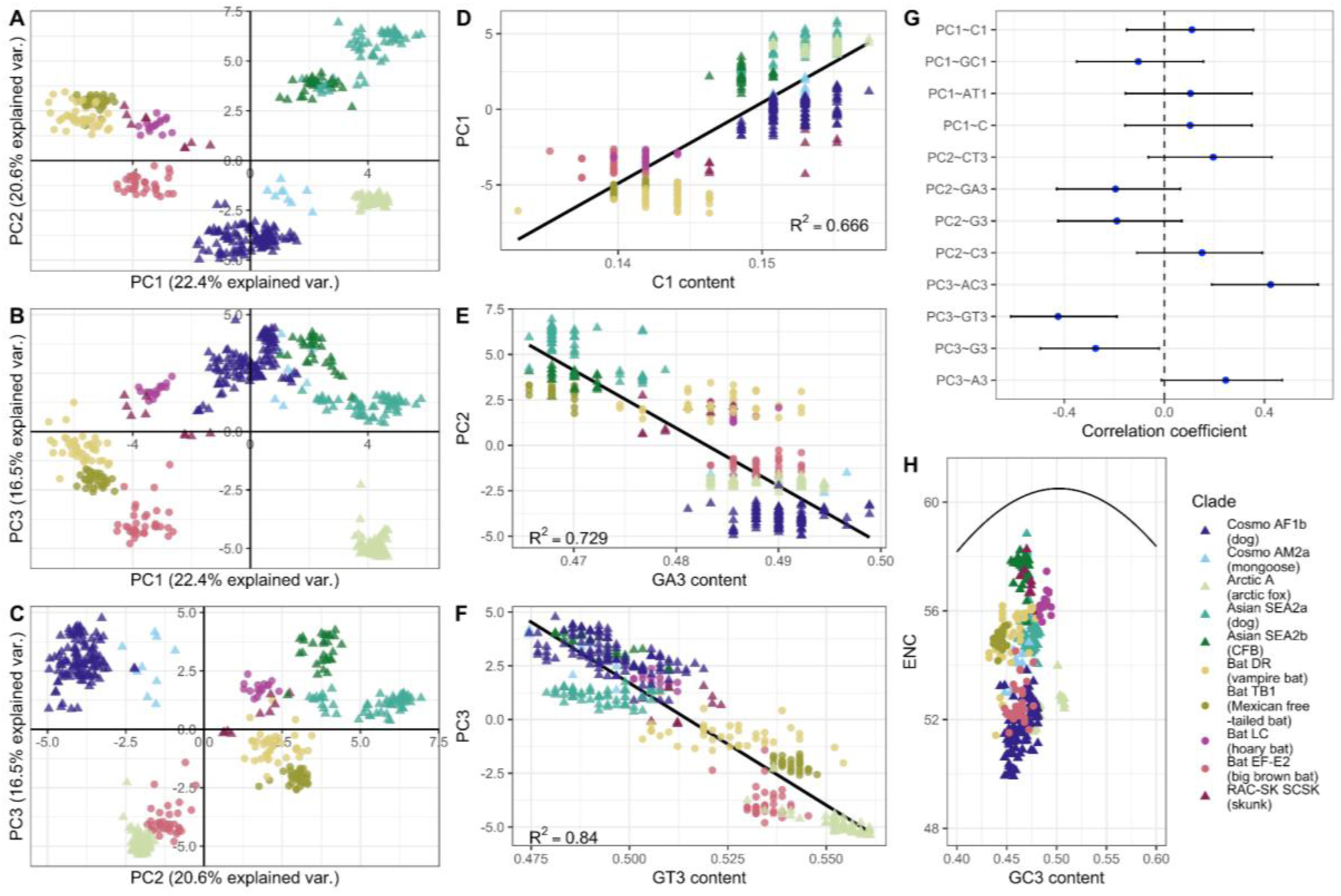
Codon usage between species-specific clades is different, and shaped by purine and UpA content. (A-C) Principal component analysis of raw codon usage values. **(D)** Cytosine content at the third position is correlated with PC1. **(E)** Purine content at the third position is negatively correlated with PC2. **(F)** Guanine and thymine content at the third position is negatively correlated with PC3. **(G)** The correlation coefficients of the 4 most highly correlated nucleotide composition metrics for individual codons with the loadings of each principal component. Error bars represent the 95% confidence interval. **(H)** An ENC-GC3 plot showing that the points do not follow the curve expected if codon bias was due purely to GC content. The point colour represents the RABV clade. Point shape represents whether the host species is a bat (round) or a carnivore (triangle). CFB = Chinese ferret badger.

To investigate whether RABV codon usage patterns more closely resembled those of the clades’ host species than its non-host species, we calculated the normalised codon adaptation index for each clade with *Canis familiaris* (domestic dog), *Vulpes lagopus* (Arctic fox), *Desmodus rotundus* (vampire bat) and *Eptesicus fuscus* (big brown bat) host codon usage values as references (Figure 4A). We did not find evidence of substantial differences in nCAI across hosts in any of the RABV clades, and therefore no evidence that RABV N gene codon usage biases mimic those of their host species. The bat-associated RABV clades, along with the skunk-associated clade RAC-SK SCSK, had higher nCAI values than the carnivore-associated clades across all four host species; only the Bat EF-E2 clade, however, had a mean nCAI above 1 for all host species. nCAI was strongly correlated with UpA content (adjusted R^2^ = 0.596 (95%CI 0.572 - 0.620); Figure 4B), and significantly so when phylogenetic structure was taken into account (pMCMC < 0.001), implying that the higher nCAI values observed across hosts in the bat-associated clades may be due to their low UpA content, as mammalian genes are also UpA deficient (Karlin & Mrázek 1997). CpG content was also negatively correlated with nCAI, but to a slightly lesser extent (adjusted R^2^ = 0.553 (0.525 - 0.581); pMCMC < 0.001).

**Figure 4:**
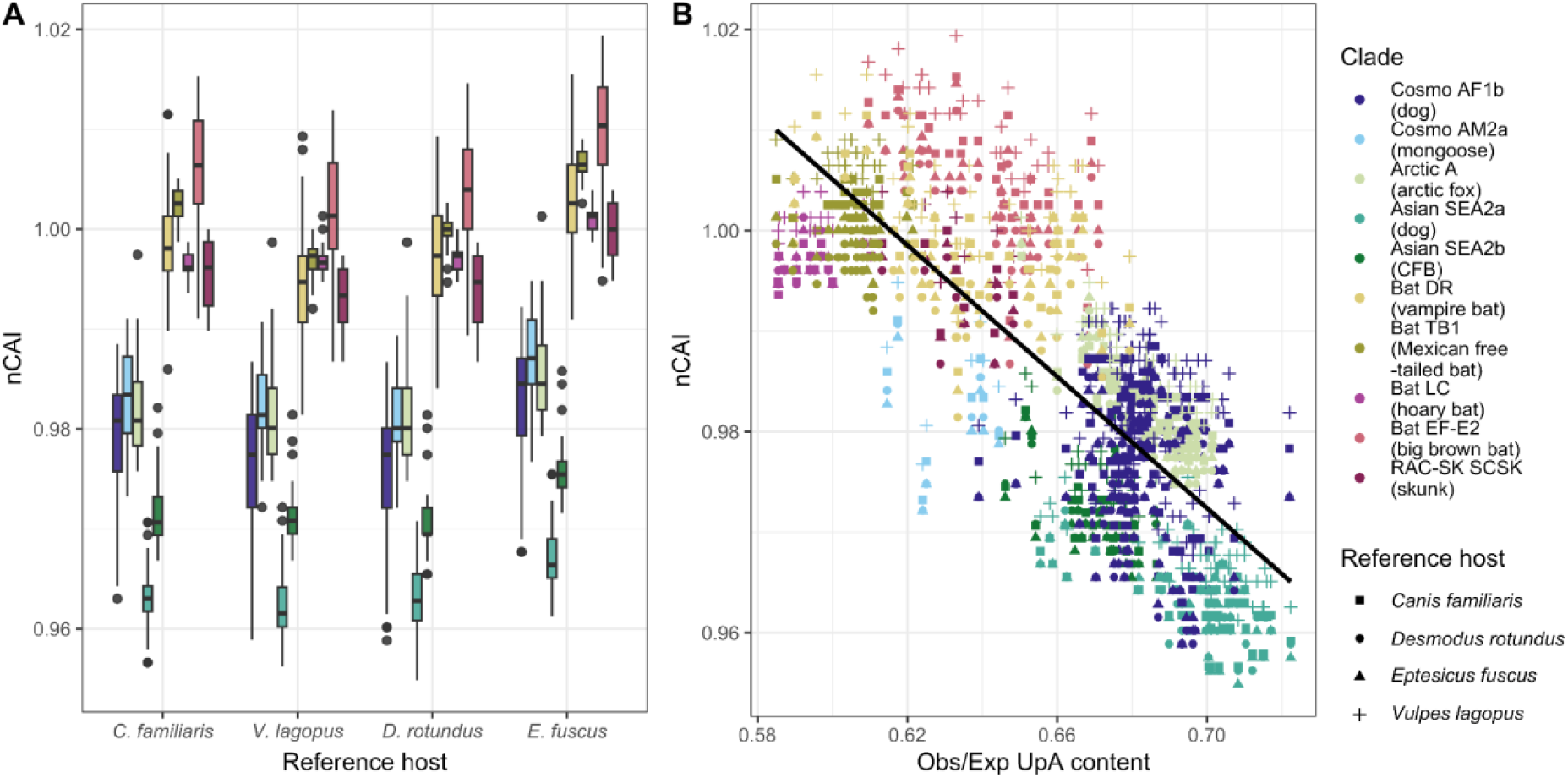
Normalised codon adaptation index (nCAI) values are higher in bat-associated RABV clades than in carnivore-associated clades; this is likely driven by UpA content. **(A)** Normalised codon adaptation index (nCAI) of each host-specific RABV clade. *Canis familiaris* (domestic dog), *Vulpes lagopus* (arctic fox), *Desmodus rotundus* (vampire bat) and *Eptesicus fuscus* (big brown bat) codon usage values are used as reference sets; bar colour represents RABV minor clade. **(B)** nCAI is negatively correlated with UpA content. Point colour represents the host-species-specific clade, point shape represents the reference host dataset. CFB = Chinese ferret badger.

### Dinucleotide content

As CpG and UpA were shown to have a large influence over RABV nCAI, and are known to be selected against in RNA viruses due to pressure from immune mechanisms such as ZAP (Meagher *et al*. 2019; Odon *et al*. 2022), we further investigated the differences in these dinucleotides between the RABV clades. CpG and UpA underrepresentation (i.e., a ratio of observed-to-expected CpG or UpA content less than 1) was seen across all host-species-specific minor clades (Figure 5). Bat-associated clades had a lower CpG and UpA content than carnivore-associated clades (mean observed-to-expected CpG content of 0.569 in carnivore-associated clades *versus* 0.488 in bat-associated clades; mean UpA of 0.682 in carnivore-associated clades *versus* 0.625 in bat associated clades). The Bat TB1 and Bat LC clades in particular displayed high levels of CpG suppression, whereas Bat DR and Bat EF-E2 have intermediate CpG and UpA contents, similar to that of the skunk clade RAC-SK SCSK. On the other hand, the two dog-associated clades (Cosmopolitan AF1b and Asian SEA2a), as well as the CFB-associated clade Asian SEA2b, showed the least evidence of CpG suppression. A similar pattern emerges when comparing UpA content across the clades, but with less of a clear distinction between the bat- and carnivore-associated clades than as seen for CpG. The differences in CpG and UpA content between bat- and carnivore-associated clades was not found to be significant after accounting for phylogenetic structure, however (pMCMC = 0.102 for CpG content; pMCMC = 0.068 for UpA content), and a very large proportion of the variation in CpG and UpA content could be explained by phylogenetic relatedness (0.985 (95% highest posterior density (HPD): 0.935 - 0.980) for CpG content; 0.965 (95%HPD: 0.941 - 0.983) for UpA content).

**Figure 5:**
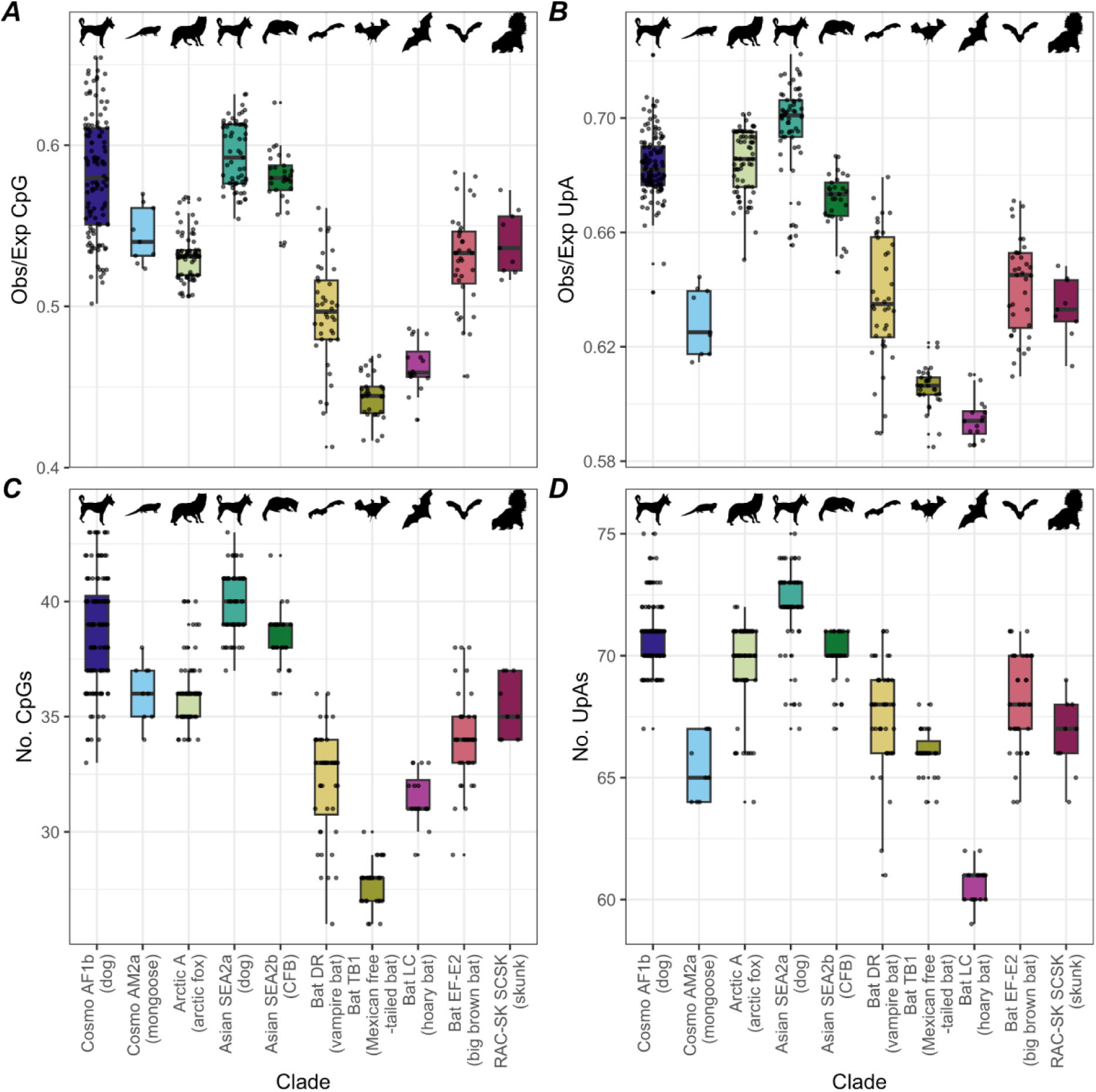
Ratio of observed-to-expected CpG and UpA content and CpG and UpA counts across host-species-specific minor clades. **(A)** Ratio of observed-to-expected CpG content, where a value of 1 indicates no bias, >1 indicates overrepresentation of CpG, and <1 indicates underrepresentation. **(B)** Ratio of observed-to-expected UpA content. **(C)** Number of CpG dinucleotides by clade. **(D)** Number of UpA dinucleotides by clade.

To investigate the evolutionary history of CpG and UpA content in the rabies virus, we used ancestral sequence reconstruction and calculated the CpG and UpA content of the resulting ancestral RABV sequences. This analysis estimates that the common ancestor of all the clades (pre-bat-to-carnivore shift) had an observed-to-expected CpG content ratio similar to that of the modern bat-associated clades (0.484 (95%CI 0.400 - 0.567) at the RABV common ancestor node, compared to a mean of 0.488 in the modern bat-associated clade sequences), implying that low CpG may have been the ancestral state (Figure 6). Similar CpG content is estimated at the common ancestor nodes of the bat-associated clades (0.476; 95%CI 0.418 - 0.556) and the carnivore associated clades (0.493; 95%CI 0.420 - 0.583), which may imply that high CpG content in carnivore-associated clades emerged after the main bat-to-carnivore host shift; the confidence intervals for the ancestral CpG content are wide enough to encompass the mean CpG contents for both the modern bat- and carnivore-associated clades, however. Ancestral UpA content may have also more closely resembled the content found in modern bat-associated clades, with the RABV common ancestor node being estimated to have an observed-to-expected UpA ratio of 0.637 (95%CI 0.585 - 0.689) compared to a mean of 0.625 in modern bat-associated clades and 0.682 in carnivore-associated clades. The common ancestor of the carnivore clades was estimated as having a UpA content of 0.644 (0.587 - 0.701), and the common ancestor of the bat clades as having a ratio of 0.621 (0.577 - 0.659), but as with the ancestral CpG content, the confidence intervals are very broad.

**Figure 6:**
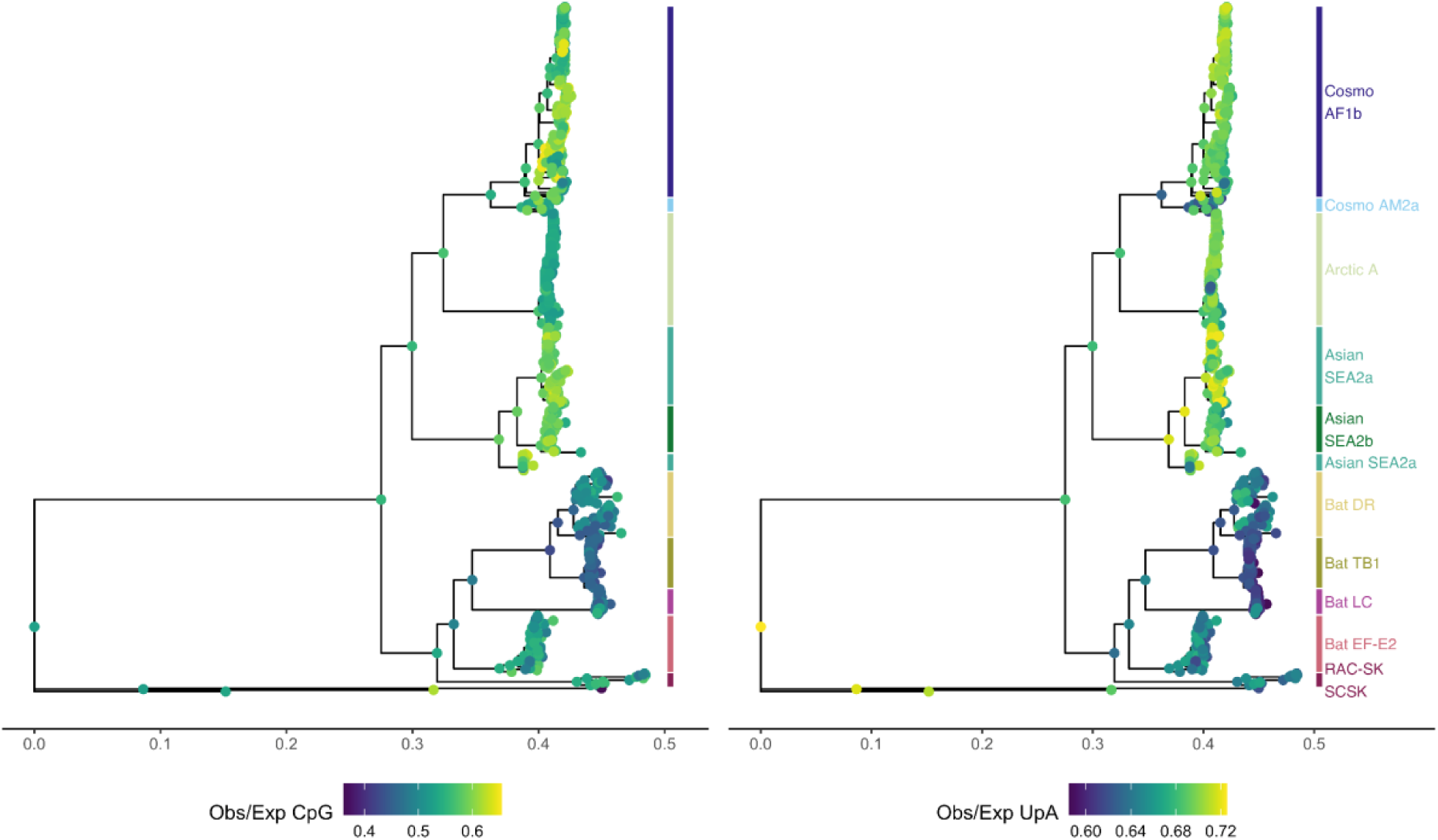
Low CpG and UpA content appears to be the ancestral state of RABV. Phylogenetic tree of all the N gene sequences with ancestral node sequences reconstructed and CpG (left) and UpA (right) content calculated. Point colour represents the observed dinucleotide content as a proportion of the expected dinucleotide content given the sequence’s nucleotide content.

When comparing the frequencies of mouse-ZAP’s optimal binding motifs in the N gene sequences to the estimated number of motifs given the sequences’ nucleotide and dinucleotide contents, we found that carnivore-associated RABV clade N gene sequences on average contained approximately the same number of ZAP binding motifs as expected (mean observed-to-expected ZAP motif ratio of 0.995), while bat-associated clades had fewer motifs than expected (mean ratio of 0.755). In particular, the Chinese ferret badger-associated Asian SEA2b clade sequences and the big brown bat-associated Bat EF-E2 clade sequences had far fewer ZAP binding motifs than expected (Figure 7; mean ratios of 0.574 and 0.465 respectively). Unlike CpG and UpA content alone, the difference in observed-to-expected ZAP motif ratios between bat- and carnivore-associated clades was found to be significant after accounting for phylogenetic structure (pMCMC = 0.025).

**Figure 7:**
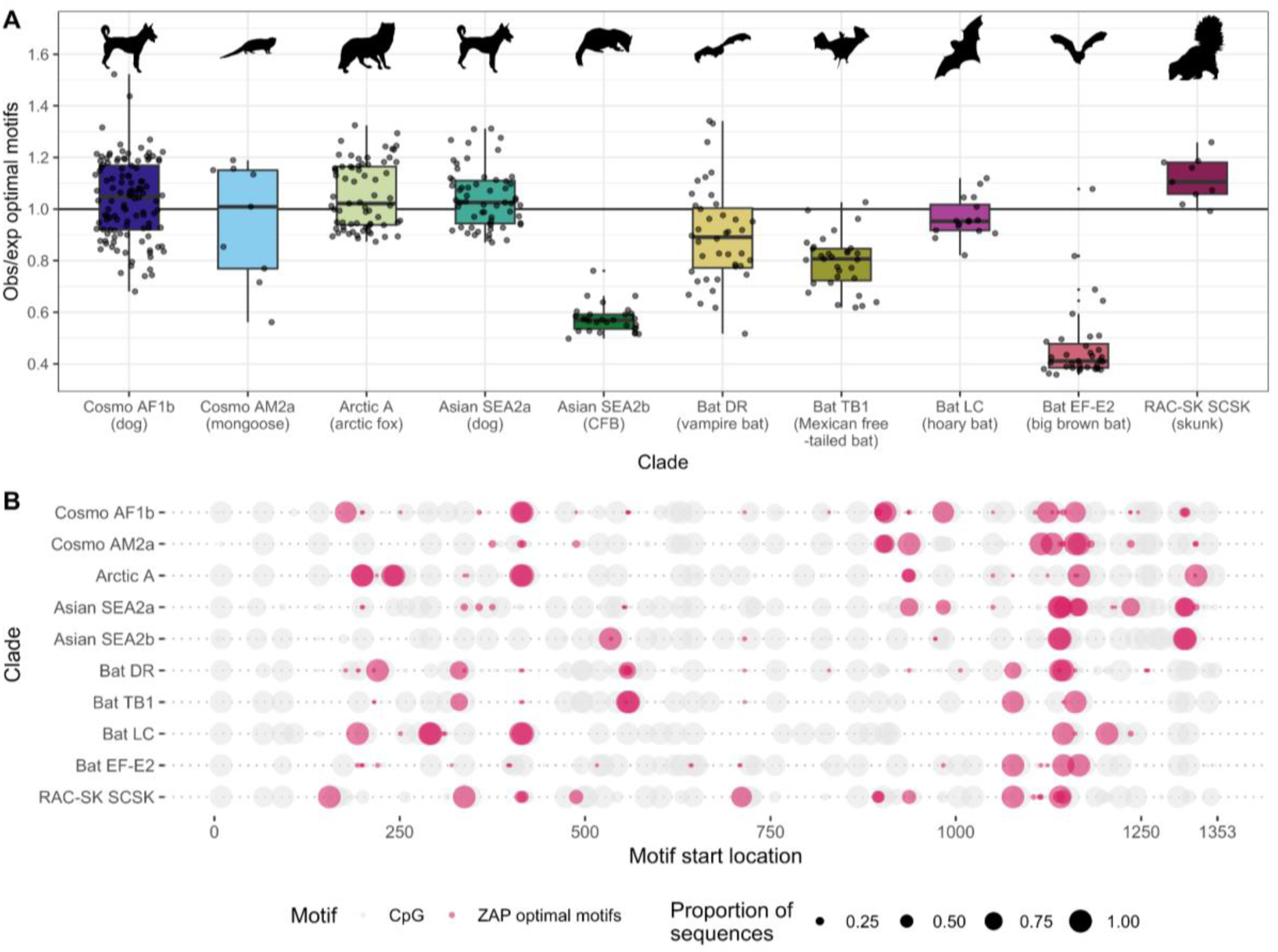
Bat-associated RABV N gene sequences contain on average fewer ZAP binding motifs than would be expected based on their nucleotide and dinucleotide contents. **(A)** Ratio between the number of observed and expected ZAP binding motifs by species-specific RABV clade. **(B)** Locations of CpG dinucleotides (grey) and ZAP binding motifs (pink) on the N gene. Point size represents the proportion of sequences within each clade where each motif is present at each locus.

## Discussion

We found that much of the variation in codon usage between host-species-specific rabies virus clades can be explained by differences in C1, GA3 and GT3 content. We also found that CpG content is lower in the bat-associated RABV clades than in carnivore-associated clades. The strong under-representation of ZAP optimal binding motifs in the Asian SEA2b and Bat EF-E2 clades could indicate that host ZAP exerts a higher selection pressure on some RABV clades than others, but further study would be required to confirm this.

The results of our principal component analysis indicated that most host-species-specific rabies virus clades had distinct patterns of codon usage, whereas the RAC-SK south-central skunk and Bat LC clades clustered closely together, despite belonging to different major clades and infecting hosts belonging to different taxonomic families. While the bat- and carnivore-associated clades appear to broadly split across PC1, the two dog-associated clades are the most distant clades across PC2.

Further analysis revealed that PC2 is significantly correlated with purine content at the third codon position (GA3), which explains 73% of the variation along PC2. Higher purine content in the virus’s genomic RNA, which would correspond to a lower purine (or higher pyrimidine) content in the coding RNA sequences used in our analysis, is associated with restricted access of polymerase to the genomic RNA through the nucleocapsid, and thus reduced transcription (Gumpper *et al*. 2019). It is currently unclear whether the modulation of transcription or another factor drives the differences in purine content at the third codon position between RABV clades; as the two dog-associated clades have very different purine contents, it seems unlikely that these differences are purely due to host adaptation. As little is currently known about the role of purine and pyrimidine content in negative-strand RNA viruses, further work on the topic could reveal the evolutionary forces driving the differences between these clades. While the correlation of both the PC3 values and PC3 loadings with guanine and thymine content at the third codon position are strong, the biological implications of varying GT3 content are even less well understood than purine content. Guanine and thymine are referred to as “keto” nucleotides due to a shared keto group in their molecular structure, but we could not find any previous research on how the prevalence of this pair affects viral fitness. The significant relationship between the PC3 loadings and the GT3 content of individual codons suggests that preference for or against GT3 occurs at the codon level, rather than being due to differences in amino acid content.

PC1’s correlation with cytosine content at the first codon position may be related to the avoidance of CpG dinucleotides, but if this had a strong influence over codon usage we would also expect to see a stronger correlation with GC3 content; synonymous substitutions to cytosine at position 1 can only result in a new CpG dinucleotide within an arginine codon, and all of the clades investigated strongly preferred the AGA codon for arginine, but synonymous substitutions to guanine or cytosine at position three can form new CpG dinucleotides within or across codons. Combined with little correlation being found between the PC1 loadings and the C1 content of individual codons, it may be more likely that the correlation between the PC1 values and C1 content is due to differences in the prevalence of C-beginning amino acids between sequences.

CpG content in RNA viruses is almost universally lower than would be expected based on GC content due to selective pressure from the host’s zinc-finger antiviral protein (ZAP) which specifically targets motifs containing CpG dinucleotides (Takata *et al*. 2017). UpA, another target of ZAP, and is also depleted in RNA virus genomes, but to a lesser extent than CpG (Odon *et al*. 2022). Bat-associated RABV clades have a CpG content that would be considered typical for negative-strand RNA viruses, while the carnivore-associated clades have noticeably higher CpG content, though still within the known range for mammalian negative-strand RNA viruses (Simmonds *et al*. 2013). A similar, but less clear, pattern is observed for UpA. Statistically significant differences in CpG and UpA content between bat- and carnivore-associated clade groups were not found when accounting for phylogenetic structure, but to some extent this is unsurprising; differences in nucleotide composition between sequences will directly influence their phylogenetic distance. Carnivore host genes generally do not appear to have a higher CpG content than bat genes (Shaw *et al*. 2021), so the difference in CpG content between bat- and carnivore-associated clades is not likely to be due to host mimicry. This finding is also opposite to a similar analysis conducted in coronaviruses, where coronaviruses with bat hosts had a higher CpG content than coronaviruses found in other mammals (MacLean *et al*. 2021; Nchioua *et al*. 2020). Ancestral sequence reconstruction suggests that low CpG content may have been the ancestral state of the rabies virus, but as the confidence interval on this value is very wide and the substitution model used is not dinucleotide-aware, this result should be treated with caution.

We also found that on average, bat-associated RABV clades had fewer CpG-based ZAP optimal binding motifs (in the format C(n_m_)G(n)CG, as identified in mice) than would be expected in randomised sequences maintaining identical nucleotide and dinucleotide content to the RABV sequences, whereas most of the carnivore-associated clades had approximately as many ZAP-optimal motifs as would be expected. This may support the possibility of weaker selection pressure against CpG and the mouse-derived ZAP binding motif in most carnivore-associated clades compared to the bat-associated clades, and a strong selection pressure against the ZAP binding motif in the Bat EF-E2 and Asian SEA2b clades. One possibility is that ZAP efficacy varies between the host species; Nchioua et al. found that the lentiviruses that infected primate species that had the less effective “extralong” isoform of ZAP had higher CpG content than the lentiviruses infecting primate species with the more effective “long” isoform (Nchioua *et al*. 2025). Another is that the rabies virus may be more exposed to ZAP within bat hosts than within carnivore hosts; there is much ongoing discussion as to the differences between the immune systems of bats and other mammals. Despite having very low observed-to-expected ZAP motif ratios, Bat EF-E2 had the highest CpG content of the bat-associated clades, and Asian SEA2b had a CpG content comparable to the other carnivore-associated clades, which may suggest that the mouse-derived ZAP motif may simply more closely match the ZAP motif that the host ZAP actually targets in these clades than in the other clades. Further research into ZAP binding motifs in species other than mice may shed some light on this.

The CAI analysis suggests that RABV codon usage does not match the codon usage of specific hosts. Our findings instead suggest that bat-associated RABV codon usage more closely resembles both carnivore- and bat-hosts’ codon usage than carnivore-associated RABV clades do, but that the codon usage of each RABV clade is not more similar to either bat or carnivore codon usage. Codon usage tables were also not available for all host species so not every clade’s codon could be directly compared to that of their associated host, but no evidence of host-specific codon usage mimicking was found for the direct comparisons that could be made. It seems likely that the higher CAI values observed in the bat-associated RABV clades are simply reflective of their lower CpG and UpA content, as vertebrate genes also have both low CpG content due to the deamination of methylated CpG sites (Sved & Bird 1990) and UpA content (Karlin & Mrázek 1997). As CoCoPUTs’ codon usage data for non-human animals cannot be filtered for genes associated with certain tissue types, it is possible that the CAI analysis could be affected by the high tissue-specificity of codon usage in multicellular organisms (Payne & Alvarez-Ponce 2019; Plotkin *et al*. 2004), especially if some host species’ codon usage data are biased towards certain tissue types.

One limitation of our investigation is that we focused only on ten minor RABV clades with clear species associations, leaving some major clades unrepresented. Different clades have different sampling time ranges, but these differences are small compared to the estimated divergence times of the lineages, and are therefore assumed to not affect our findings. Another is that we analysed only the N gene; as whole-genome sequencing is heavily biased towards dog-associated clades, many of the clades included in our analyses had few or no whole-genome sequences available. The N gene was chosen as it was the most commonly sequenced gene, providing the largest number of sequences across the clades, but other genes may be subject to different selective pressures. Further investigation to confirm whether the patterns observed in our study are consistent across the genome may become feasible as whole-genome RABV sequencing becomes more widespread, and should also be investigated in other lyssaviruses.

We found differences in the codon usage between host-species-specific RABV clades, a large proportion of which appear to be driven by non-GC nucleotide composition. CpG content and observed-to-expected ZAP motif ratios were also higher in carnivore-associated clades than in bat-associated clades, which may suggest differences in selection pressure from ZAP between the groups, and should be explored further. These results may also have implications for the efficacy of codon-deoptimised or CpG-enriched live attenuated vaccines across different host-species-specific clades should they be developed for the rabies virus, as they have been for Influenza A (Sharp *et al*. 2023).

## Acknowledgements

The authors wish to thank Denise Marston, Liam Brierley, Spyros Lytras, Hollie French and Jocelyn Perez-Lazo for their helpful discussions and feedback. This work was supported by the EPSRC DTP (EP/T517896/1 to RD); the Canadian Institutes of Health Research; and Wellcome (224670/Z/21/Z to KH and 218518/Z/19/Z to MA).

## Author contributions

RD: Conceptualisation, data curation, formal analysis, investigation, methodology, visualisation, writing - original draft, writing - review and editing

JD: Supervision, methodology, writing - review and editing

MA: Writing - review and editing

CC: Supervision, writing - review and editing

KH: Supervision, funding acquisition, writing - review and editing

## Data availability

All of the code and data used in this study are available in the GitHub repository https://github.com/RowanDurrant/rabies-codon-bias. The accession numbers of all sequences used are available in Supplementary Table S1.

## Supplementary material

**Supplementary Table S1:** Accession numbers for the sequences used in our analysis by host species.

| Clade (Host) | Accession numbers |
| --- | --- |
| Asian SEA2b (Chinese ferret badger) | FJ598135, FJ712195, FJ712196, FJ719751, FJ719753, FJ719755, GU647092, HQ118114, JN974877, JQ950446, JQ950448, KF726852, KP319185, KP319186, KP319187, KP319188, KP319191, KP319193, KP319194, KP319195, KP319199, KP319201, KP319203, KP319204, KP319205, KP319206, KP319207, MT068612, MT068615 |
| Asian SEA2a (dog) | DQ666289, DQ666290, DQ866105, DQ866106, DQ866108, DQ866109, DQ866110, DQ866112, DQ866116, DQ866119, EU086182, EU086183, EU828653, FJ561726, FJ561727, FJ561728, FJ866829, FJ866830, FJ866831, GQ472468, GQ472470, GQ472473, GQ472477, GU358653, HM486355, HM486356, HM486368, HM486369, HM486374, HM486379, HM486381, JF819605, JF819615, JN974823, JN974824, JN974828, JN974829, JN974838, JN974842, JN974845, JN974870, JN974871, JN974872, JN974873, JQ970480, KT221107, KT894558, KT894572, KT894573, KT894577, KT894578, KX148264, KY451767, MG201879, MG201880, MG201884, MG201919, MG201921, MN784132 |
| Arctic A (Arctic fox) | AY352488, AY352501, JN258588, JN258592, JX987747, KU198460, KU198462, KU198464, KU198466, KU198467, KU198472, KU198476, KU198477, KU198478, KX036361, KX036362, KX036364, KX036365, KX036366, KX954123, LT598537, LT598538, LT598539, LT598540, LT598541, LT598543, LT909550, MG458313, MN233885, MN233886, MN233892, MN233896, MN233900, MN233948, MN233949, MN233950, MN233952, MN233953, MN233955, MN233956, MN233958, MN233960, MN233963, MN233967, MN233968, MN233969, MN233973, MN233974, MN233975, MN233977, MN233978, MN233982, MN233985, MN233988, MN233991, MN233992, MN233998, MN234000, MN234002, MN234004, MN234008, MN234016, MN234017, MN234018, MN234023, MN234043, MN234046, MN234055, MN384717 |
| Cosmopolitan AF1b (dog) | AB284509, AB284511, AB284512, AB284513, AB284514, EU853582, EU853584, EU853586, EU853587, EU853590, HM179504, HM179505, HM179506, KF155002, KR534217, KR534218, KR534219, KR534228, KR534229, KR534230, KR534233, KR534238, KR534244, KR534250, KR534251, KR534252, KR534253, KR906735, KR906736, KR906739, KR906741, KR906742, KR906744, KR906745, KR906746, KR906747, KR906748, KR906749, KR906751, KR906756, KR906757, KR906759, KR906762, KR906763, KR906765, KR906766, KR906767, KR906770, KR906773, KR906780, KR906784, KR906790, KR906792, KT336432, KT336433, KT336434, KT336435, KT336436, KT336437, KX148203, KX148204, KX148206, KX148208, KY210224, KY210243, KY210247, KY210249, KY210253, KY210263, KY210264, KY210279, KY210287, KY210304, KY563717, LC029890, MN726804, MN726826, MN726827, MT454631, MT454635, |
|  | MT454637, MT454641, MT454643, MT454644, MT454645, OK317992, OK317993, OK317995, OK317996, OK317998, OK317999, OP375406, OP375407, OP375410, OP375413, OP375414, OP375415, OP375418, OP375421, OP375423, OR045939, OR045949, OR270973, OR270979, OR270997, OR270998, OR270999, OR271001, OR271003, PV855388, PV855390, PV855392, PV855393, PV855394, PV855397, PV855415 |
| Bat LC (hoary bat) | AF351845, AF351846, AF351858, AF394883, AF394884, GU644712, GU644713, GU644714, GU644715, GU644716, GU644717, GU644718, GU644719, GU644720, GU644721, JQ685947 |
| Cosmopolitan AM2a (mongoose) | AY854505, AY854525, AY854531, AY854544, AY854546, AY854558, AY854567, AY854573, AY854576 |
| RAC-SK SCSK (skunk) | EU345002, EU345003, EU345004, JQ685938, JQ685968, MW055084, MW055085, MW055086, MW055087 |
| Bat DR (vampire bat) | AB519642, AF070449, AF351847, AF351852, AY854587, AY854592, AY854594, AY877433, AY877434, AY877435, GU592648, KF656696, KF864234, KF864322, KF864397, KM594040, KM594041, KM594042, KP202393, KT023101, MN968374, MN968375, MN968377, MN968386, MT891038, MW249020, MW579786, MW579820, MW579824, MW579833, PQ379829, PQ379831, PQ379832, PQ379834, PQ379836, PQ379849, PQ510120, PQ603086, PQ671599, PQ671601 |
| Bat EF-E2 (big brown bat) | AF351828, AF351831, AF351832, AF351833, AF351855, AF351861, AF351862, AY039227, AY039228, AY039229, GU644652, GU644654, GU644655, GU644656, GU644657, GU644658, GU644659, GU644660, GU644661, GU644662, GU644663, GU644664, GU644665, GU644666, GU644667, GU644668, GU644669, GU644670, GU644671, GU644676, GU644677, GU644684, GU644689, GU644690, GU644695 |
| Bat TB1 (Mexican free-tailed bat) | AF351849, AF394876, GU644760, GU644761, GU644762, GU644763, GU644764, GU644765, GU644766, GU644767, GU644768, GU644769, GU644770, GU644771, GU644772, GU644773, GU644774, GU644775, GU644776, GU644777, GU644778, GU644779, GU644780, GU644781, GU644782, GU644784, GU644785, GU644786, GU644787, GU644788, JQ685905 |

**Supplementary Table S2:** Mean RSCU values across different clades. Green highlight indicates that the codon is significantly underrepresented (RSCU value < 0.6), orange highlight indicates that the codon is significantly overrepresented (RSCU value > 1.6). Bold indicates the most preferred codon for each amino acid in each clade.

| Amino acid | Codon | Asian SEA2b (CFB) | Cosmo AF1b (dog) | Asian SEA2a (dog) | Arctic A (Arctic fox) | Cosmo AM2a (mongoose) | RAC-SK SCSK (skunk) | Bat LC (hoary bat) | Bat EF-E2 (big brown bat) | Bat TB1 (Mexican free tailed bat) | Bat DR (vampire bat) |
| --- | --- | --- | --- | --- | --- | --- | --- | --- | --- | --- | --- |
| Phe | UUU | 0.874 | 0.635 | 0.822 | 0.802 | 0.642 | **1.004** | 0.972 | **1.067** | **1.123** | 0.897 |
|  | UUC | **1.126** | **1.365** | **1.178** | **1.198** | **1.358** | 0.996 | **1.028** | 0.933 | 0.877 | **1.103** |
| Leu | UUA | 0.986 | 0.558 | 1.092 | 0.449 | 0.723 | 0.602 | 0.903 | 0.966 | 1.045 | 1.012 |
|  | UUG | **1.252** | 1.245 | 1.037 | 1.679 | **1.505** | **1.728** | **2.114** | **1.798** | **1.637** | **1.642** |
|  | CUU | 0.889 | 0.356 | 1.000 | 0.603 | 0.332 | 0.940 | 0.859 | 0.604 | 0.868 | 0.691 |
|  | CUC | 0.718 | 0.542 | 0.513 | 0.521 | 0.704 | 0.678 | 0.528 | 0.708 | 0.520 | 0.661 |
|  | CUA | 1.074 | **1.697** | 1.044 | 0.944 | 1.290 | 0.735 | 0.704 | 0.522 | 0.503 | 0.584 |
|  | CUG | 1.080 | 1.603 | **1.315** | **1.803** | 1.446 | 1.318 | 0.891 | 1.401 | 1.427 | 1.411 |
| Ile | AUU | **1.145** | 0.977 | **1.112** | **1.07** | **1.069** | 0.921 | 1.033 | 0.854 | 1.039 | **1.100** |
|  | AUC | 0.735 | 0.927 | 0.846 | 0.929 | 0.915 | 0.862 | 0.858 | 1.033 | 1.039 | 0.855 |
|  | AUA | 1.120 | **1.096** | 1.042 | 1.001 | 1.016 | **1.217** | **1.109** | **1.113** | 0.920 | 1.045 |
| Val | GUU | 1.179 | 0.997 | 1.018 | 1.089 | 0.989 | 1.011 | **1.280** | 1.068 | **1.224** | **1.192** |
|  | GUC | **1.333** | **1.454** | **1.381** | **1.139** | **1.458** | 1.073 | 1.004 | 0.738 | 0.932 | 0.954 |
|  | GUA | 0.655 | 0.563 | 0.943 | 0.881 | 0.548 | 0.622 | 0.711 | 0.749 | 0.634 | 0.728 |
|  | GUG | 0.833 | 0.985 | 0.658 | 0.892 | 1.005 | **1.295** | 1.004 | **1.444** | 1.210 | 1.126 |
| Ser | UCU | 1.222 | 1.286 | **1.557** | 1.222 | 1.181 | 1.276 | 1.407 | 1.699 | **1.729** | **1.579** |
|  | UCC | 1.232 | 0.793 | 1.000 | 0.842 | 0.895 | **1.535** | 0.640 | 0.702 | 0.783 | 0.776 |
|  | UCA | **1.604** | **1.846** | 1.272 | **1.733** | **1.886** | 1.274 | **1.741** | **1.818** | 1.581 | 1.535 |
|  | UCG | 0.486 | 0.735 | 0.588 | 0.658 | 0.667 | 0.607 | 0.305 | 0.389 | 0.320 | 0.445 |
|  | AGU | 0.973 | 0.981 | 1.057 | 0.842 | 1.048 | 0.645 | 1.269 | 0.895 | 0.955 | 1.019 |
|  | AGC | 0.481 | 0.359 | 0.527 | 0.703 | 0.324 | 0.663 | 0.640 | 0.497 | 0.631 | 0.646 |
| Pro | CCU | **1.241** | **1.560** | 1.047 | **1.652** | 1.417 | 1.020 | **1.632** | **1.615** | **1.766** | **1.939** |
|  | CCC | 1.000 | 0.928 | **1.202** | 1.172 | 1.083 | 0.856 | 0.735 | 0.721 | 0.984 | 0.601 |
|  | CCA | 1.017 | 0.891 | 0.945 | 1.168 | **1.472** | 0.936 | 0.941 | 0.539 | 0.492 | 0.513 |
|  | CCG | 0.741 | 0.620 | 0.806 | 0.007 | 0.028 | **1.188** | 0.691 | 1.125 | 0.758 | 0.948 |
| Thr | ACU | **1.398** | 1.141 | **1.509** | **1.539** | 1.353 | 1.358 | 0.978 | 0.964 | 1.417 | 1.307 |
|  | ACC | 0.954 | 1.243 | 0.879 | 0.721 | 0.837 | 0.981 | 1.368 | 1.353 | 0.777 | 0.838 |
|  | ACA | 1.198 | **1.463** | 1.026 | 0.690 | **1.489** | **1.460** | **1.515** | **1.415** | **1.546** | **1.622** |
|  | ACG | 0.450 | 0.153 | 0.586 | 1.051 | 0.320 | 0.199 | 0.139 | 0.268 | 0.258 | 0.233 |
| Ala | GCU | 1.322 | 1.000 | **1.507** | 1.348 | 1.058 | 1.482 | 0.852 | 1.129 | **1.484** | 1.389 |
|  | GCC | 1.027 | 1.078 | 0.794 | 0.879 | 0.990 | 0.725 | 0.982 | 0.990 | 0.883 | 0.805 |
|  | GCA | **1.503** | **1.710** | 1.494 | **1.634** | **1.695** | **1.705** | **1.804** | **1.836** | 1.153 | **1.413** |
|  | GCG | 0.148 | 0.211 | 0.205 | 0.139 | 0.257 | 0.088 | 0.363 | 0.046 | 0.480 | 0.393 |
| Tyr | UAU | **1.074** | **1.249** | **1.144** | **1.505** | **1.280** | **1.266** | 0.851 | **1.408** | **1.330** | **1.198** |
|  | UAC | 0.926 | 0.751 | 0.856 | 0.495 | 0.720 | 0.734 | **1.149** | 0.592 | 0.670 | 0.802 |
| His | CAU | 0.531 | **1.219** | 0.780 | 0.923 | **1.265** | **1.083** | **1.231** | **1.229** | **1.308** | **1.311** |
|  | CAC | **1.469** | 0.781 | **1.220** | **1.077** | 0.735 | 0.917 | 0.769 | 0.771 | 0.692 | 0.689 |
| Gln | CAA | **1.193** | **1.023** | **1.569** | **1.200** | 1.000 | 0.674 | **1.013** | **1.058** | **1.119** | 0.830 |
|  | CAG | 0.807 | 0.977 | 0.431 | 0.800 | 1.000 | **1.326** | 0.988 | 0.942 | 0.881 | **1.170** |
| Asn | AAU | 0.945 | **1.069** | 0.970 | **1.172** | 0.883 | **1.088** | 0.918 | **1.327** | **1.236** | **1.274** |
|  | AAC | **1.055** | 0.931 | **1.030** | 0.828 | **1.117** | 0.912 | **1.083** | 0.673 | 0.764 | 0.726 |
| Lys | AAA | **1.051** | 0.910 | 0.821 | 0.495 | 0.885 | 0.683 | 0.589 | 0.679 | 0.960 | 0.960 |
|  | AAG | 0.949 | **1.090** | **1.179** | **1.505** | **1.115** | **1.317** | **1.411** | **1.321** | **1.040** | **1.040** |
| Asp | GAU | **1.017** | 0.996 | 0.717 | **1.041** | 0.992 | **1.010** | 0.867 | 0.902 | 0.956 | 0.952 |
|  | GAC | 0.983 | **1.004** | **1.283** | 0.959 | **1.008** | 0.990 | **1.133** | **1.098** | **1.044** | **1.048** |
| Glu | GAA | 0.913 | 0.873 | 0.804 | 0.690 | 0.805 | **1.222** | 0.833 | 0.766 | 0.919 | 0.869 |
|  | GAG | **1.087** | **1.127** | **1.196** | **1.310** | **1.195** | 0.778 | **1.168** | **1.234** | **1.081** | **1.131** |
| Cys | UGU | **1.023** | **1.948** | 0.498 | **1.333** | **1.429** | **1.111** | **1.062** | **1.015** | **1.333** | **1.030** |
|  | UGC | 0.977 | 0.052 | 1.502 | 0.667 | 0.571 | 0.889 | 0.938 | 0.985 | 0.667 | 0.970 |
| Arg | CGU | 0.780 | 0.804 | 0.780 | 0.791 | 0.746 | 0.492 | 0.480 | 0.531 | 0.504 | 0.380 |
|  | CGC | 0 | 0.017 | 0 | 0 | 0.030 | 0.552 | 0.240 | 0.224 | 0.252 | 0.370 |
|  | CGA | 0.780 | 0.342 | 0.776 | 0.261 | 1.061 | 0.376 | 0.480 | 0.030 | 0 | 0.273 |
|  | CGG | 0.260 | 0.474 | 0.266 | 0.772 | 0.317 | 0.523 | 0.495 | 0.792 | 0.512 | 0.417 |
|  | AGA | **2.319** | **3.189** | **2.805** | **3.317** | **2.844** | **2.462** | **2.415** | **3.617** | **2.757** | **2.804** |
|  | AGG | 1.863 | 1.174 | 1.374 | 0.859 | 1.002 | 1.595 | 1.890 | 0.807 | 1.974 | 1.757 |
| Gly | GGU | 1.079 | 0.770 | 0.872 | 0.811 | 0.838 | 0.882 | 0.699 | 0.940 | 0.556 | 0.435 |
|  | GGC | 0.401 | 0.702 | 0.549 | 0.417 | 0.603 | 0.794 | 0.949 | 0.529 | 0.819 | 0.931 |
|  | GGA | 1.134 | 1.102 | 1.215 | 1.013 | 1.382 | **1.168** | 0.975 | 1.199 | **1.660** | **1.648** |
|  | GGG | **1.387** | **1.426** | **1.365** | **1.758** | **1.176** | 1.156 | **1.379** | **1.332** | 0.966 | 0.986 |

**Supplementary Table S3:** PCA loadings for each codon on PC1 and PC2.

| Amino acid | Codon | PC1 | PC2 | PC3 |
| --- | --- | --- | --- | --- |
| F | TTT | -0.1238798 | 0.14969785 | -0.1538584 |
|  | TTC | 0.12312555 | -0.1532243 | 0.15259887 |
| L | TTA | -0.0812072 | 0.22367578 | 0.02686364 |
|  | TTG | -0.1129219 | -0.0225038 | -0.1916261 |
|  | CTT | 0.02306152 | 0.26314298 | -0.0511886 |
|  | CTC | -0.1000353 | 0.04885798 | -0.0145167 |
|  | CTA | 0.1135647 | -0.1556209 | 0.20204684 |
|  | CTG | 0.07898578 | -0.1565916 | -0.1300544 |
| I | ATT | 0.08630567 | 0.06288623 | 0.01381255 |
|  | ATC | -0.0620328 | -0.1352308 | -0.1030095 |
|  | ATA | -0.0041873 | -0.1197346 | 0.12262433 |
| V | GTT | -0.0542682 | 0.09606773 | -0.0303476 |
|  | GTC | 0.15332263 | -0.0518635 | 0.22602061 |
|  | GTA | 0.1380224 | 0.1369166 | -0.1354166 |
|  | GTG | -0.1946201 | -0.0896962 | -0.07509 |
| S | TCT | -0.1495211 | 0.15423219 | -0.0620752 |
|  | TCC | 0.08025414 | 0.14053837 | 0.07468375 |
|  | TCA | -0.0820463 | -0.1959373 | -0.000233 |
|  | TCG | 0.13319661 | -0.1151713 | 0.09597072 |
|  | AGT | -0.0534769 | 0.12409925 | 0.12714272 |
|  | AGC | -0.0260652 | 0.1144503 | -0.1953197 |
| P | CCT | -0.1396412 | -0.1177935 | -0.1435618 |
|  | CCC | 0.20641153 | 0.02744473 | -0.0499682 |
|  | CCA | 0.19515143 | -0.045743 | 0.01343401 |
|  | CCG | -0.1603017 | 0.09819678 | 0.06693523 |
| T | ACT | 0.10407946 | 0.11476606 | -0.14015 |
|  | ACC | -0.0981148 | -0.1134341 | 0.10882896 |
|  | ACA | -0.2371434 | -0.0057607 | 0.120301 |
|  | ACG | 0.18184363 | 0.02786802 | -0.2112205 |
| A | GCT | 0.05003959 | 0.18662366 | -0.1271572 |
|  | GCC | -0.0089584 | -0.1729656 | 0.12979821 |
|  | GCA | 0.05665423 | -0.1914721 | 0.04631538 |
|  | GCG | -0.1365341 | 0.06016717 | 0.04883212 |
| Y | TAT | 0.0406744 | -0.132285 | -0.2318439 |
|  | TAC | -0.041297 | 0.13122295 | 0.23190373 |
| H | CAT | -0.2084364 | -0.1216455 | -0.0120494 |
|  | CAC | 0.20005047 | 0.14040413 | 0.00407316 |
| Q | CAA | 0.20443162 | 0.13159382 | 0.00098762 |
|  | CAG | -0.1674602 | -0.1455563 | 0.03870188 |
| N | AAT | -0.1227503 | -0.0262692 | -0.2027536 |
|  | AAC | 0.14256955 | 0.0729617 | 0.19341221 |
| K | AAA | -0.0862157 | 0.03246694 | 0.23386082 |
|  | AAG | 0.10029931 | -0.0907991 | -0.204241 |
| D | GAT | -0.0507114 | -0.1465419 | 0.00189217 |
|  | GAC | 0.02860627 | 0.19559701 | 0.05931421 |
| E | GAA | -0.1028884 | 0.00177118 | 0.17134389 |
|  | GAG | 0.14081741 | -0.0689321 | -0.1470548 |
| C | TGT | -0.0306328 | -0.2471806 | 0.11614228 |
|  | TGC | 0.03588002 | 0.24888704 | -0.0826329 |
| R | CGT | 0.22584217 | -0.0709505 | 0.07567385 |
|  | CGC | -0.2191445 | 0.06700388 | -0.0692887 |
|  | CGA | 0.14547106 | 0.12879737 | 0.17176766 |
|  | CGG | -0.0233685 | -0.1253969 | -0.2416757 |
|  | AGA | 0.00143952 | -0.1466299 | -0.1945023 |
|  | AGG | -0.1182627 | 0.17147498 | 0.10522635 |
| G | GGT | 0.15645291 | -0.0134029 | 0.00694933 |
|  | GGC | -0.1959211 | -0.0051093 | 0.1054936 |
|  | GGA | -0.1587444 | 0.09814174 | -0.0294782 |
|  | GGG | 0.20129191 | -0.1132681 | -0.0777299 |

**Supplementary Figure S1:**
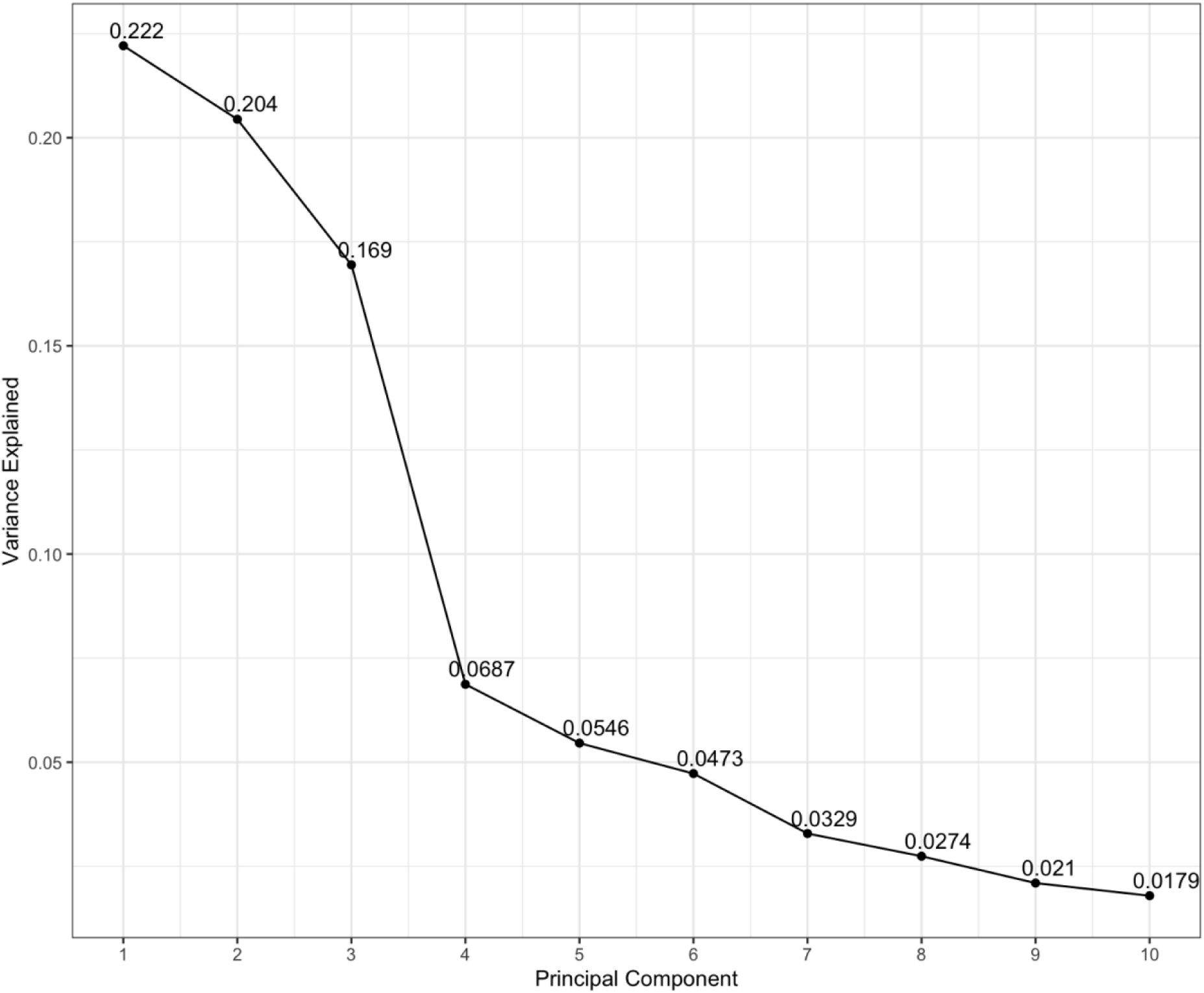
Scree plot for the PCA of raw codon usage values.

**Supplementary Figure 2:**
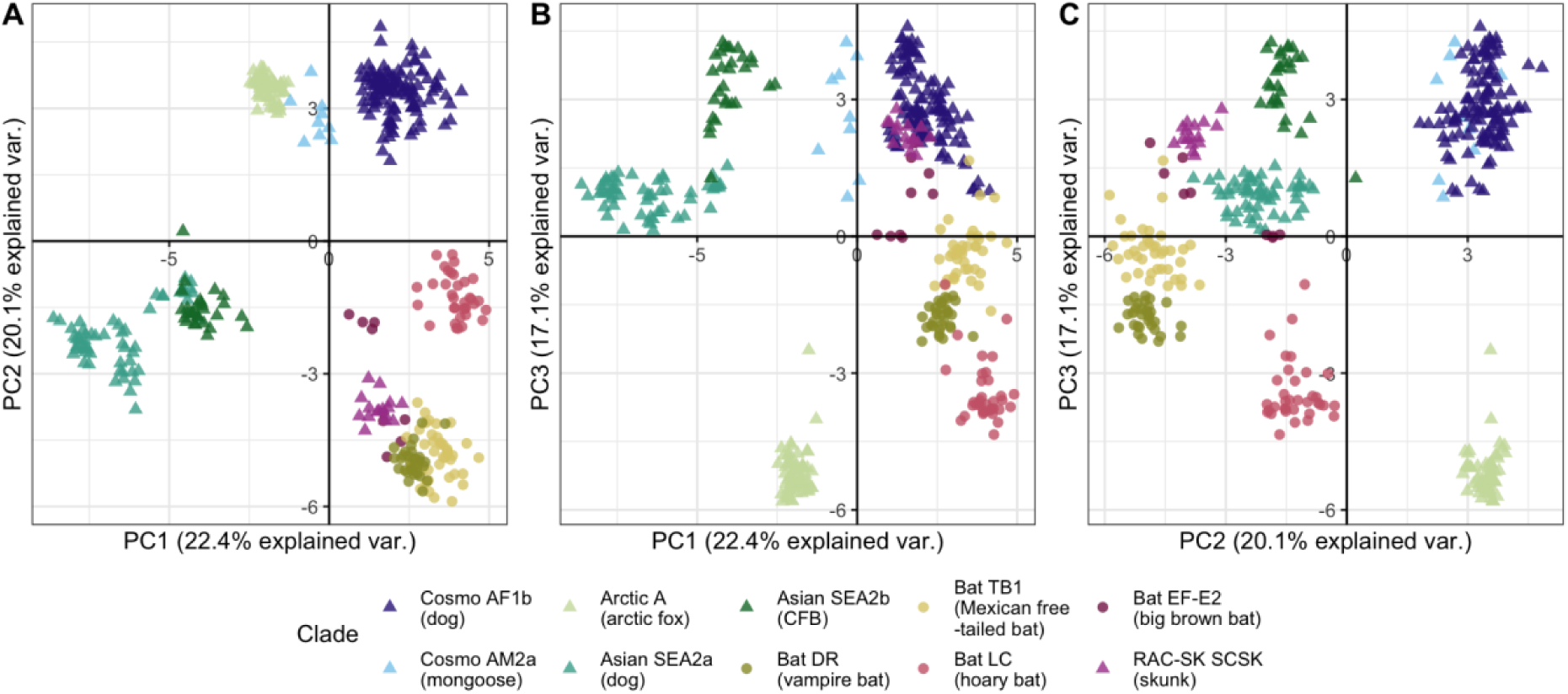
PCA using RSCU values.

